# Phosphorylation-mediated regulation of the essential splicing factor PUF60

**DOI:** 10.64898/2026.05.17.725788

**Authors:** Mohammed S. Ali, Paul L. Boutz

**Affiliations:** Department of Biochemistry and Biophysics, University of Rochester School of Medicine and Dentistry, Rochester, NY 14642, USA; Center for RNA Biology, Rochester, NY 14642, USA; Wilmot Cancer Institute, Rochester, NY 14642, USA

## Abstract

PUF60 is a splicing factor related to the polypyrimidine-tract binding protein U2AF2. PUF60 is deleted in developmental disorders such as Verheij syndrome and amplified in approximately 8% of cancers. Thus, both increases and decreases in PUF60 expression can have profound physiological effects. However, little is known about how changes in PUF60 expression impact global splicing patterns. Here, we created a model system of CRISPRa/i in mouse stem cells (mESCs) to transcriptionally upregulate or downregulate Puf60. Our results uncovered extensive transcriptional, post-transcriptional, and post-translational regulation of Puf60 protein expression. We observed that Puf60 protein levels in normal mESCs drop dramatically at a critical cell density, leading to cell death. Puf60 is very essential in stem cells, and its repression causes cell death and impacts specific splicing events, including its own splicing autoregulation, providing valuable insights into the functional consequences of PUF60 dysregulation. Analysis of phosphoprotein data revealed phosphorylation of threonine at the N-terminus of PUF60. Our results showed that mutating threonine to glutamate downregulates the protein and alters its localization. Thus, our study reveals a novel regulatory mechanism of Puf60 phosphorylation that mediates its function and may be related to its frequent overexpression in cancer cells.

## Introduction

Alternative splicing is increasingly recognized as a hallmark of cancer progression. Through alternative splicing, a single mRNA can produce multiple isoforms that drive oncogenic processes, serve as tumor markers, and confer resistance to therapeutics [1]. Dysregulated splicing can affect many genes and signaling pathways involved in tumor development and progression. Splicing factor mutations have been identified in a wide range of solid and hematological tumors [2, 3]. These observations underscore the importance of understanding how splicing regulatory networks contribute to tumor initiation, progression, and treatment resistance.

Pre-mRNA splicing is a critical step in post-transcriptional gene regulation and is mediated through coordinated interactions between cis-acting RNA elements and trans-acting splicing factors that direct spliceosome assembly at the appropriate 5′ and 3′ splice sites. This process removes introns and joins exons to generate mature mRNA transcripts. Most human genes undergo alternative splicing (AS), greatly expanding transcriptomic and proteomic diversity by generating multiple RNA isoforms from a single gene [4]. Importantly, splice-site selection is not uniformly regulated, as distinct subsets of introns exhibit differential dependence on specific splicing factors[5]. Consequently, dysregulation of AS resulting from mutations, altered expression, amplification, or deletion of splicing regulatory factors and cis-acting elements has been implicated in numerous human diseases, including cancer [6-8].

Poly U binding splicing factor 60 (PUF60) is a splicing factor involved in the recognition of the polypyrimidine tract and assembly of the U2 small nuclear ribonucleoprotein (U2 snRNP) complex during the early stages of spliceosome formation. Through interactions with core spliceosomal components, PUF60 contributes to accurate splice-site recognition and the regulation of alternative splicing events [9]. Aberrant expression of Puf60 has been reported in multiple malignancies, including hepatocellular carcinoma, ovarian cancer, bladder cancer, renal cancer, and lung cancer [10-14]. In contrast, loss-of-function mutations or haploinsufficiency of PUF60 are associated with developmental disorders such as Verheij syndrome (8q24.3 microdeletion syndrome), which is characterized by neurological, cardiac, ocular, and renal abnormalities, as well as developmental delay [15-21]. These findings suggest that PUF60 plays context-dependent roles in both development and disease; however, the molecular mechanisms underlying these distinct physiological outcomes remain poorly understood.

Here, we established a CRISPRa/i system to investigate Puf60-regulated splicing in mouse embryonic stem cells (mESCs). Our results reveal that Puf60 is tightly regulated at both the transcriptional and translational levels. We further demonstrate that a conserved threonine residue at position 64 plays a critical role in phosphorylation-dependent regulation of Puf60. Importantly, phosphorylation modulates Puf60 subcellular localization and may impact cellular metabolic activity. Collectively, our results uncover a previously unrecognized link between Puf60 phosphorylation, subcellular localization, and possibly, metabolic regulation.

## Results

### Splicing factor alterations across human cancers

Aberrant expression of splicing factors has been commonly related to disease progression [22]. Analysis of datasets from the Cancer Genome Atlas (TCGA) revealed that multiple splicing factors containing the UHM domain (U2AF homology motif), such as RBM17, RBM39, U2AF2, and PUF60, are altered in cancer patients, and a higher proportion of these alterations are amplifications (Fig. 1A). PUF60 was the most altered in these cancer patients. GISTIC analysis identified recurrent amplification of the PUF60 locus in tumor samples, and there is a correlation between PUF60 gene copy number and mRNA fold change, with a nearly two-fold increase in mRNA when there are more than two copies (Fig. 1B). DepMap data showed that PUF60 is an essential gene in more than 90% of cells, as determined by RNAi or CRISPR knockout screens (Fig. 1C). We hypothesized that modulating Puf60 expression in the same cell line could be particularly informative, since patient data are likely affected by low signal-to-noise due to genetic background variation beyond amplification or deletion itself. We therefore developed a CRISPRa/i model to upregulate and downregulate Puf60 at the transcription levels (Fig. 2D). We used a second-generation CRISPRa Suntag system and one of the most repressive KRAB domains (ZIM3) for CRISPRi [23-25]. We generated stable mouse stem cells expressing either the CRISPRa or the CRISPRi machinery, which were stably integrated into the Rosa26 safe harbor locus via homology-directed repair (HDR) (Fig. 1E). We generated multiple homozygous clones confirmed by genotyping (Fig. S1A), and stable cells were evaluated for machinery expression at the protein level. All CRISPRi clones showed detectable expression, whereas only a limited number of CRISPRa clones showed detectable machinery expression (Fig. S1B).

**Fig 1.**
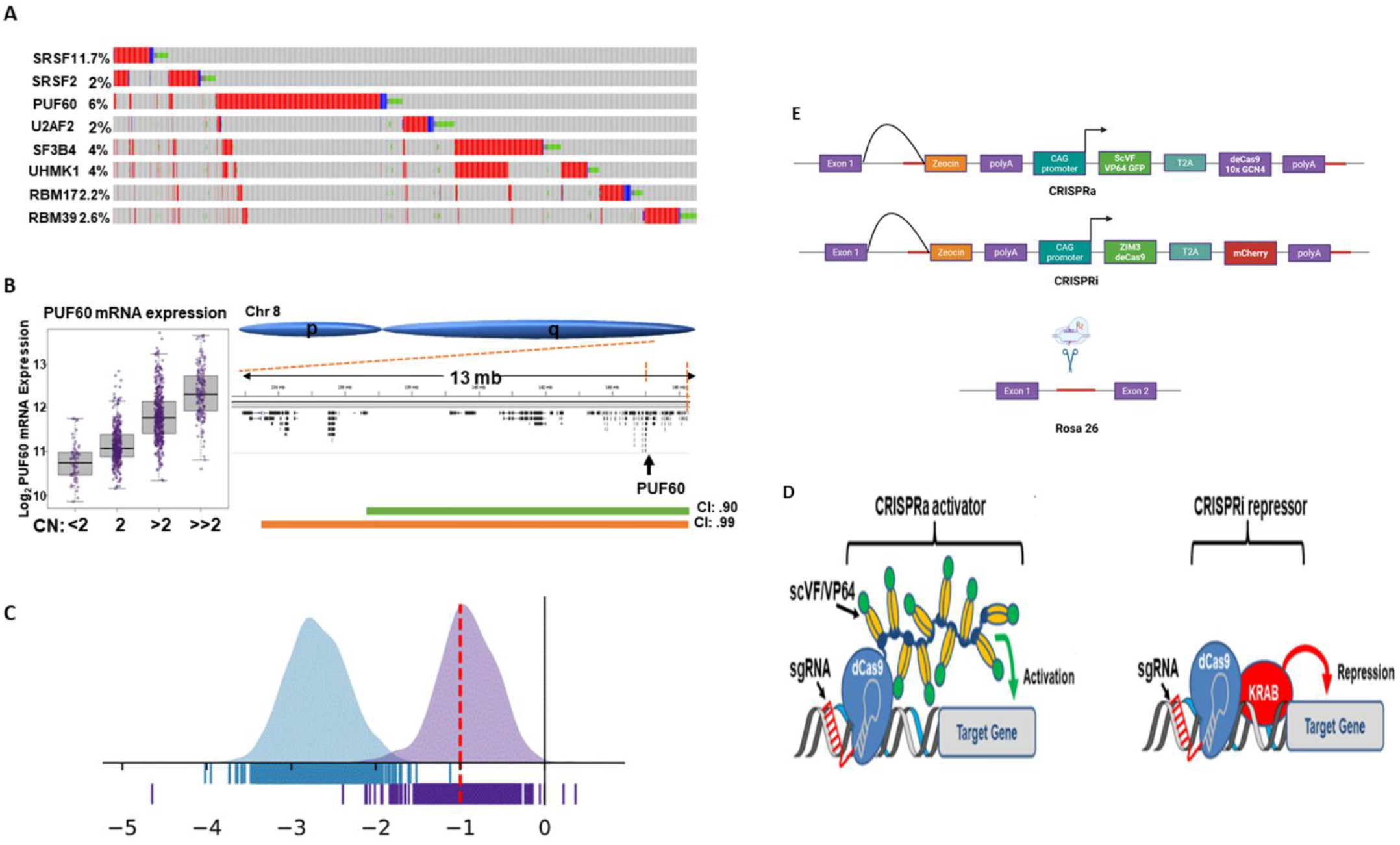
Frequent Amplification and Essentiality of Puf60 in Cancer. **A)** Pan-cancer oncoplot derived from TCGA/cBioPortal datasets. Each column represents an individual patient, with red indicating amplification, blue indicating deletion, and green indicating mutation events. **B)** Association between Puf60 copy number amplification and normalized Puf60 mRNA expression in TCGA breast cancer samples. **C)** DepMap dependency analysis demonstrates that Puf60 is essential in more than 90% of cell lines in both CRISPR knockout (blue) and RNAi knockdown (pink) datasets. A dependency score of 0 indicates a non-essential gene, whereas a score of −1 represents the median score of common essential genes. DepMap dependency analysis demonstrates that Puf60 is essential in more than 90% of cell lines in both CRISPR knockout (blue) and RNAi knockdown (pink) datasets. A dependency score of 0 indicates a non-essential gene, whereas a score of −1 represents the median score of common essential genes. **D)** Schematic overview of the CRISPRa/i system, including the second-generation SunTag CRISPRa platform and CRISPRi mediated by the Zim3 repression domain. **E)** Generation of CRISPRa and CRISPRi systems through integration of the complete machinery into the Rosa26 safe harbor locus.

**Fig 2.**
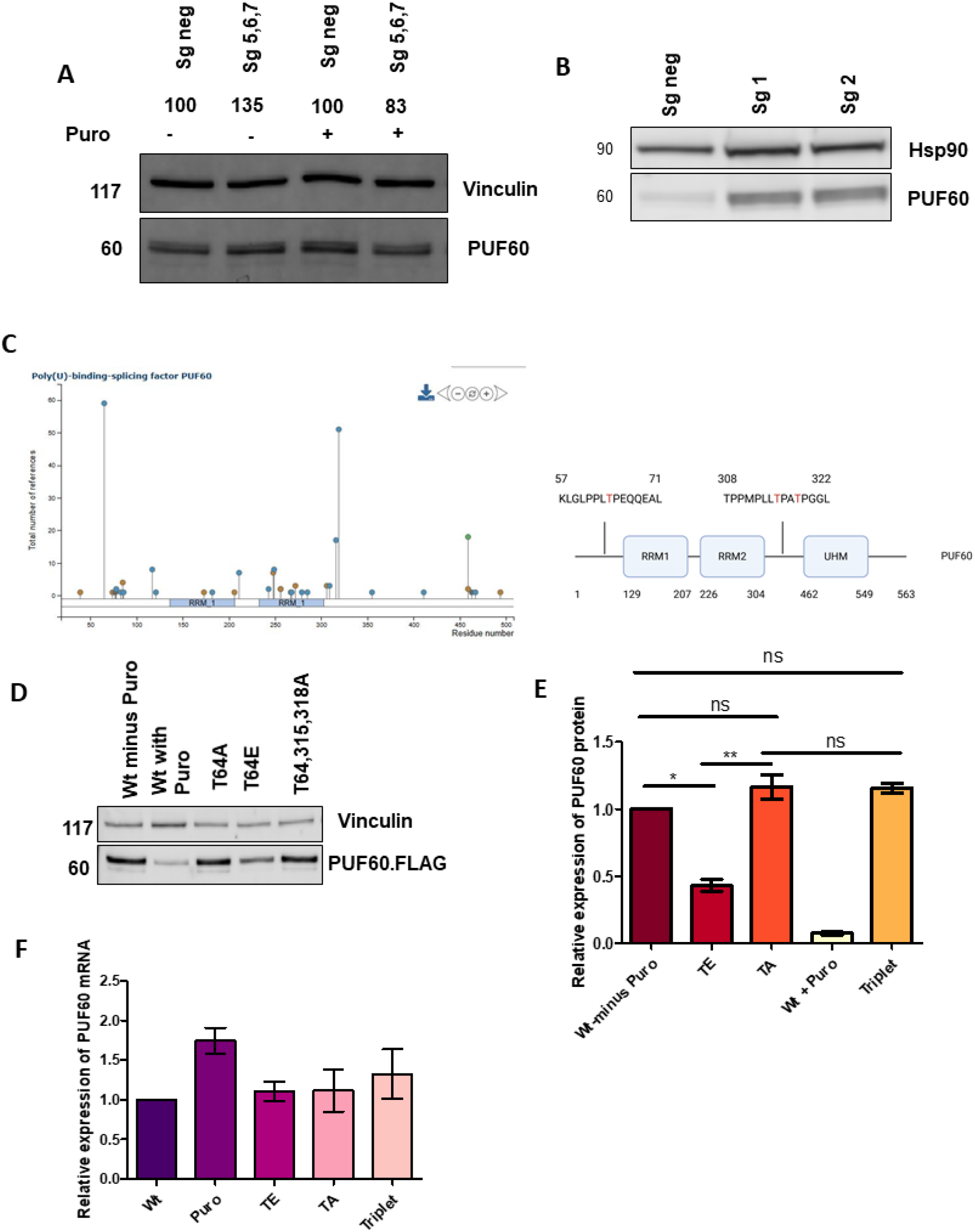
PUF60 upregulation Using CRISPRa and cDNA Overexpression Systems. **A)** Western blot analysis of CRISPRa cells transduced with control sgRNA or Puf60-targeting sgRNA after 48 hours. **B)** Western blot analysis of CRISPRa cells transduced with control sgRNA or Puf60-targeting sgRNA after 5 days. **C)** Phosphoproteomic analysis of the Puf60 gene showing post-translational modification sites. **D)** Western blot analysis of cells expressing different Puf60 cDNA constructs. **E)** Quantification of protein expression levels among the different Puf60 constructs. **F)** RNA expression quantification among the different Puf60 constructs.

### CRISPRa reveals post-transcriptional, tight regulatory control of Puf60 in mESCs

Lentiviral transduction of CRISPRa-stable mESCs with pCRISPRia-v2 carrying an Puf60-targeting sgRNA did not significantly alter Puf60 protein expression levels (Fig S1C). We validated the functionality of CRISPRa mESCs by confirming induction of genes known to be repressed in mouse stem cells, such as Dppa3. Only a few clones showed a modest increase in Dppa3 (Fig S1D) [26]. Previous studies showed that the mouse sgRNA promoter we used was less efficient in mouse stem cells than in human stem cells [27]. We used the pX330-U6-Chimeric_BB-CbH-hSpCas9 plasmid instead, as after removing the Cas9 piece and performing a transient transfection, we observed very strong Dpaa3 overexpression, which increased further when two sgRNAs were used, demonstrating a synergistic effect (Fig S1E). Although previous studies showed that the second-generation CRISPRa system effectively boosts expression of well-expressed genes by 2-fold or even higher, we tested a variety of sgRNA combinations targeting Puf60 and observed only 30-40% upregulation with one sgRNAs combination (Fig S2A; Fig. 2A) [28, 29]. When the cell line was grown for ∼ 10 days, we observed that CRISPRa cells maintained Puf60 protein expression at normal levels, whereas control cells exhibited a rapid and significant decrease at the late time point (Fig 2B). This suggested that CRISPRa is capable of maintaining RNA transcripts sufficient to express Puf60 protein at the level of proliferating cells, but it cannot exclude the possibility that Puf60 is regulated at the post-transcriptional, translational, or post-translational level. These data suggested that Puf60 protein expression is tightly controlled.

Phosphoproteomic analysis identified several previously uncharacterized phosphorylated threonine residues within the N-terminal and central regions of Puf60, including one site located upstream of the RRM1 domain and two sites located between the RRM2 and UHM domains (Fig 2C). We hypothesized that Puf60 could be regulated by phosphorylation, and to test this hypothesis, we first generated a of Puf60 cDNA expression construct, and introduced mutations by site-directed mutagenesis to phospho-dead (T64A, T315,318A) or phospho-mimetic (T64E). First, we performed a titration experiment to assess the relationship between RNA and protein expression levels of the wild-type cDNA. Increasing amounts of cDNA were transfected in a two-fold dilution series while maintaining a constant total DNA concentration by supplementing with GFP-expressing plasmid. This strategy controlled for variability in Lipofectamine complex formation and transfection efficiency. We made two notable observations in these experiments. First, stepwise increases in cDNA expression resulted in approximately a 2-fold increase in RNA levels, but without a proportional increase in protein expression; however, once expression surpassed a certain threshold, protein levels increased substantially (Fig. S2B, C). Additionally, the TE mutant displayed markedly lower protein expression than the other constructs, despite comparable RNA levels, consistent with altered protein stability or translation regulation (Fig. 2D– F). Together, these findings suggest that threonine 64 plays an important role in regulating Puf60 protein expression, supporting a phosphorylation-dependent regulatory mechanism.

### Mass Spectrometry Reveals Distinct Protein Interactions Associated with Mutant Puf60

To explore this further, we next determined whether these mutations affected protein-protein interactions. We performed quantitative mass spectrometry analysis of the mESCs cell line V6.5 transiently expressing several different Flag-tagged Puf60 constructs: WT, T64E, T64A, and T64,315,318A. After co-immunoprecipitation with anti-Flag antibody and with GFP expressing cells as the negative control, protein was analyzed by LC-MS/MS followed by enrichment analysis (Fig. 3A). Unsupervised hierarchical clustering of the top 200 interacting proteins revealed distinct interaction profiles among the different constructs, with the T64E mutation forming a separate cluster, while WT and other constructs formed similar clusters (Fig. 3B). Differential expression analysis identified several proteins in T64E that were not detected in other constructs, including those involved in mitochondrial (Vdac1, Vdac2, Pgm1, and Tomm40) and ubiquitin-related pathways (Ube2k, UBE2E1, and Rnf40) (Fig. 3C). KEGG enrichment analysis of proteins altered in T64E revealed significant enrichment of pathways associated with protein ubiquitination and the pentose phosphate pathway, whereas WT showed enrichment mainly in the spliceosome (Fig. 3D, E). Together, these findings demonstrate that T64E exhibits a specific proteomic signature, suggesting a potential role in regulating Pu60 function, and that the interacting protein ensemble switches from a predominantly nuclear, splicing related set in wild-type to a mostly cytoplasmic set of interactors with enrichment in metabolic and ubiquitination gene ontology classes.

**Fig 3.**
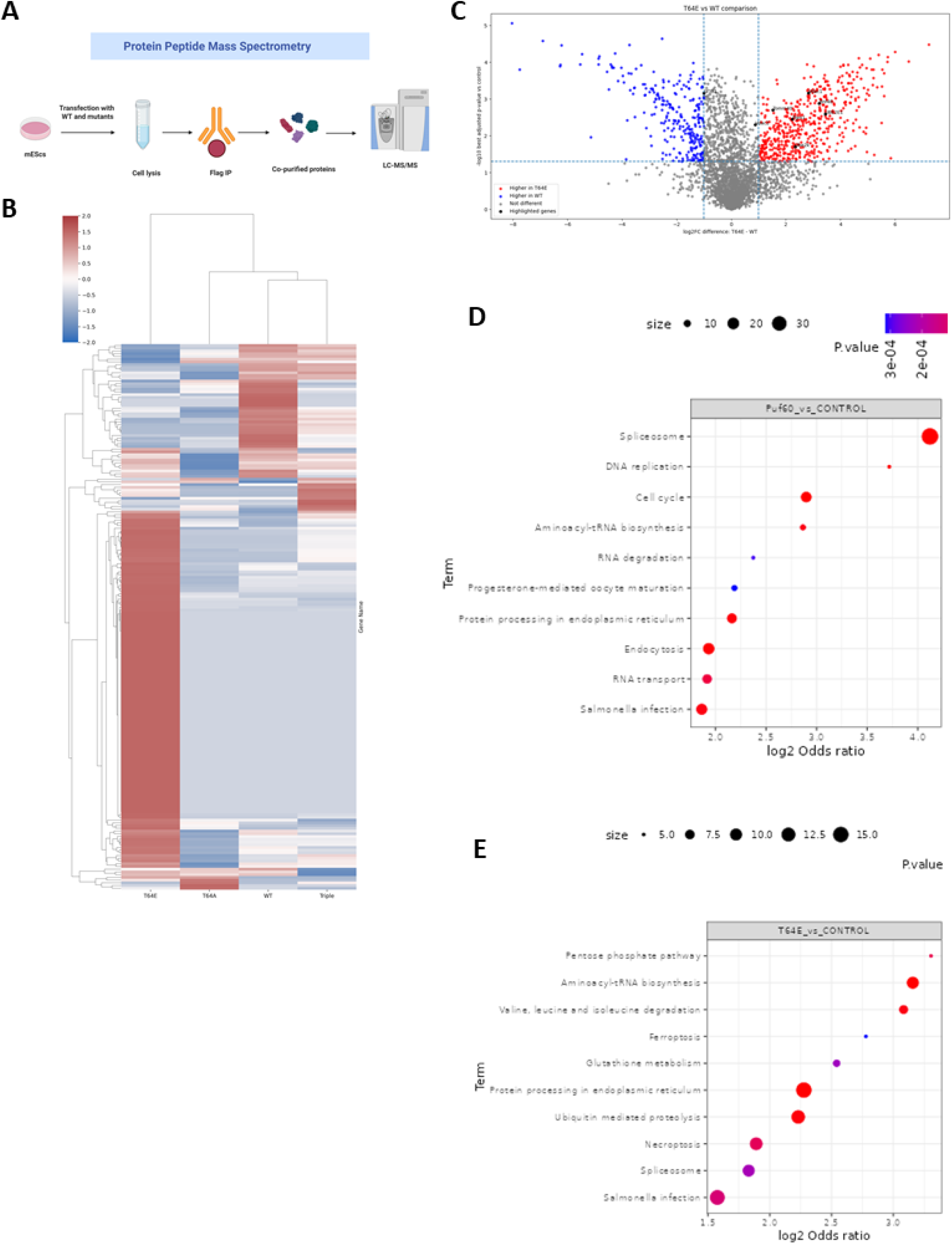
Mutation-Induced Reprogramming of the Puf60 Protein Interaction Network. **A)** Schematic Overview of the Puf60 Interactome Mass Spectrometry Workflow **B)** Clustering Analysis of Wild-Type and Mutant Puf60 Interactomes **C)** Volcano plot showing differentially regulated genes between wild-type and T64E conditions. Genes upregulated in the wild-type condition are highlighted in red, whereas genes upregulated in the T64E mutant condition are shown in blue. Selected genes of interest are labeled in black. **D)** Functional Enrichment Analysis of Wt Puf60-Associated Proteins **E)** Functional Enrichment Analysis of T64E Puf60-Associated Proteins

### Challenges in efficient repression of Puf60 expression underscore its essentiality

Results from DepMap studies have suggested that PUF60 is an essential gene in many cancer cells. To further address this possibility, we first validated the functionality of the CRISPRi mESC cell line by demonstrating efficient repression of *Oct4* expression. A single sgRNA achieved approximately 90% repression of gene expression (Table1) [30] . Based on this efficiency, one validated clone was selected for subsequent *Puf60* repression experiments. Both individual and combinatorial sgRNA approaches were evaluated; however, neither resulted in detectable changes in *Puf60* expression (Fig. S2D). We evaluated siRNA-mediated repression of Puf60 as a control approach across multiple time points; however, no significant changes in Puf60 expression were observed, indicating ineffective knockdown (Fig. S2E). However, siRNA treatment alone reduced gene expression in stem cells compared with sgNeg CRISPRi and wild-type mESC controls (Fig. S2F). Upon puromycin selection of CRISPRi-targeted cells, we achieved efficient repression of *Puf60* within 48 hours, resulting in greater than two-fold reduction in expression. Beyond 48 hours, more than 90% of the cells died and exhibited minimal media consumption, highlighting the essential role of *Puf60* in cell survival and viability (Fig. 4A, B). Consequently, subsequent experiments were limited to the 48-hour time point (Fig. 4C, D). These observations suggest that *Puf60* expression is tightly regulated and may be maintained through robust compensatory regulatory mechanisms.

**Table 1.** Validation of CRISPRi Clones Repression Efficiency Using Positive Control Target (Oct4)

| CRISPRi clones | Relative mRNA repression % |
| --- | --- |
| 8 | 60% |
| 9 | 70% |
| 18 | 90% |
| 34 | 70% |

**Table 2.**
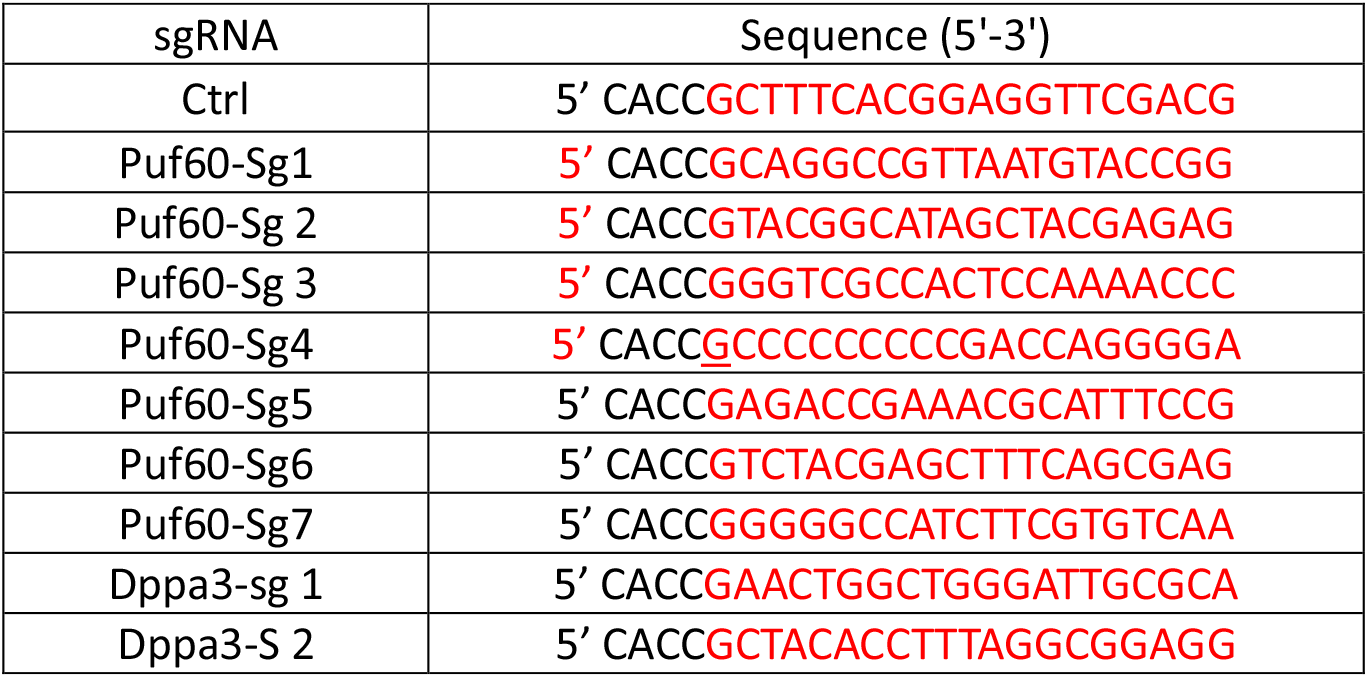
sgRNAs Used for CRISPRa-Mediated Gene Activation.

**Table 3.**
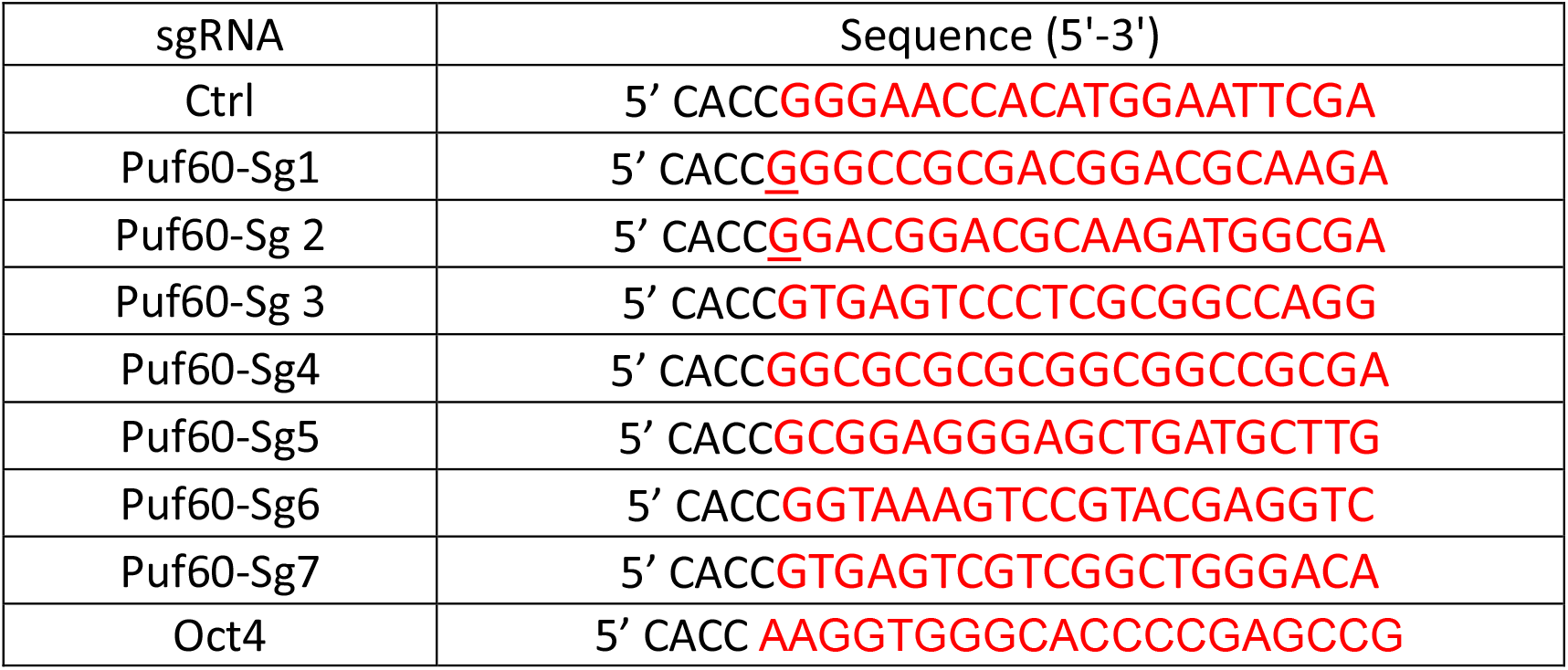
sgRNAs Used for CRISPRi-Mediated Gene Repression.

**Fig 4.**
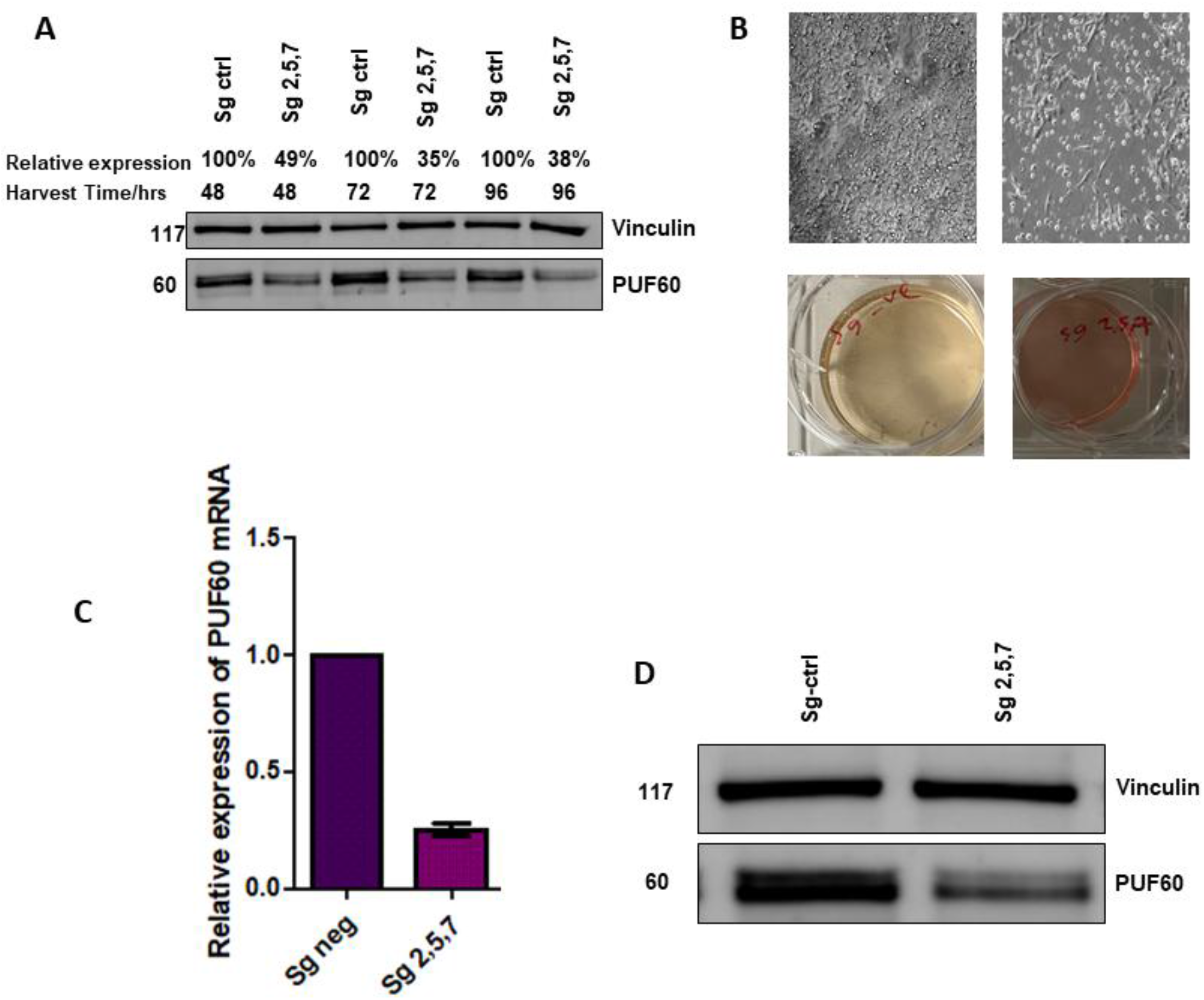
CRISPRi functional interrogation identifies Puf60 as an essential gene for cell survival. **A)** Western blot analysis of Puf60 repression by CRISPRi at multiple time points following puromycin selection. **B)** Representative images showing the cellular phenotype observed following Puf60 repression after 48 hours of CRISPRi treatment. **C)** qPCR analysis demonstrating RNA repression of Puf60 following CRISPRi targeting. **D)** Western blot validation of CRISPRi-mediated repression of Puf60.

## Discussion

UHM(U2AF-Homology Motif)-domain containing splicing factors including U2AF2 (U2AF65), RMB39 (CAPERα), RBM17 (SPF45), and PUF60 interact with other splicing factors containing a ULM (U2AF-Ligand Motif) such as U2AF1 (U2AF35) and SF3B1. The RNA binding motifs of UHM proteins typically recognize the polypyrimidine-tracts characteristic of the region between the branch site and the 3’ splice site of an intron. Thus, they link RNA sequence recognition with protein-protein interaction domains to recruit the appropriate cofactors in the determination of where a 3’ splice site will be located within a pre-mRNA. While this intricate specification process is likely essential for spliceosome function in mammalian cells, the proteins that orchestrate it are strongly susceptible to mutation in many types of cancer and pre-malignancies such as myelodysplastic syndromes. PUF60 is amplified in approximately 8% of all cancers, suggesting an oncogenic function similar to that previously established for the splicing regulator SRSF1. Loss-of-function mutations and deletions of PUF60 are causal for developmental disorders such as Verheij Syndrome. These disease associations suggest the importance of regulating PUF60 gene dosage levels precisely in physiological regimes, as dysregulation in either direction is disruptive to normal organismal function. In this study, we therefore sought to model transcriptional disturbances in the Puf60 gene using mESCs as a model system.

We first used CRISPRa to model gene amplification of Puf60. However, we were unable to boost the protein expression of Puf60 in these cell models more than a modest amount over the steady-state levels. We observed that cells undergoing stress after approximately 10 days in culture rapidly depleted Puf60 protein within 24 hours. We could maintain the steady-state amount of Puf60 with CRISPRa-mediated overexpression, suggesting that the depletion results from a transcriptional shutdown coupled with protein degradation. These data suggested that Puf60 protein levels might be regulated at the translational or post-translational level as well.

We observed in proteomic data the presence of several phosphorylated threonine residues (T64,T315, and T318) in Puf60 protein for which no known function has been attributed. To study the effects of these phosphorylation sites, we generated phospho-dead and phospho-mimetic mutants and performed co-immunoprecipitations followed by mass spectrometry to identify interacting factors. Intriguingly, the phospho-dead and wild type Puf60 proteins were largely similar in the proteins with which they interacted. However, the T64E phospho-mimetic mutant exhibited a completely different interactome. Unlike the splicing factors that the wild type Puf60 predominantly interacted with, T64E Puf60 pulled down primarily cytoplasmic factors, including mitochondrial proteins, metabolic enzymes, and ubiquitination factors. These data suggested that Puf60, upon phosphorylation at T64, transits to the cytoplasm and interacts with an entirely different set of factors.

Our data suggest that Puf60 undergoes subcellular re-localization upon phosphorylation. This re-localization likely not only results in new interactions with cytoplasmic factors that result in biological effects, but would also presumably remove Puf60 from the nucleus and its participation in splicing regulation. RNA sequencing of cells expressing the mutations is underway and will help determine whether, as we would hypothesize, Puf60 T64E mutant expressing cells will exhibit splicing patterns similar to CRISPRi or RNAi depletion of all Puf60 protein. Additionally, the altered interactions it participates in within the cytoplasm suggest that Puf60 may modulate the metabolic state of the cells. Both functions could be related to the selection of tumor cells carrying Puf60 gene amplifications. These experiments will potentially offer insight into how tumor cells dysregulate PUF60 to drive proliferative programs and may suggest novel therapeutic strategies in tumors carrying PUF60 amplifications.

## Methods

### Mammalian Cell Culture

Mouse embryonic stem cells V6.5 (mESCs) were maintained under standard culture conditions in ESC medium supplemented with fetal bovine serum and LIF at 37°C with 5% CO_2_. Cells were routinely passaged and monitored for morphology and viability.

### Lentiviral Production and Transduction

Lentiviral particles were generated in HEK293T cells by co-transfection of transfer plasmids with packaging and envelope plasmids using Lipofectamine transfection reagent. Viral supernatants were collected, filtered, and used to transduce target cells in the presence of polybrene. Transduced cells were selected using puromycin where indicated.

### Western Blot Analysis

Cells were lysed using RIPA buffer supplemented with protease and phosphatase inhibitors. Protein concentrations were quantified, and equal amounts of protein were resolved by SDS-PAGE and transferred onto PVDF membranes. Membranes were incubated with primary antibodies, followed by Cy5-conjugated secondary antibodies, and visualized by fluorescence.

### Quantitative PCR (qPCR)

Total RNA was extracted using standard RNA isolation methods and reverse-transcribed into cDNA. Quantitative PCR was performed using SYBR Green master mix on a real-time PCR system. Relative gene expression was calculated using the ΔΔCt method and normalized to housekeeping genes.

### Mass Spectrometry Analysis

Protein complexes associated with Puf60 constructs were isolated and subjected to mass spectrometry-based proteomic analysis. Samples were digested into peptides and analyzed using LC-MS/MS. Differential protein interactions and enrichment analyses were performed using standard bioinformatic pipelines.

### CRISPRa and CRISPRi Experiments

Stable CRISPRa and CRISPRi mESC lines were generated by integrating the CRISPR machinery into the Rosa26 safe harbor locus. sgRNAs targeting *Puf60* or control regions were delivered by lentiviral transduction or plasmid transfection. Gene activation or repression efficiency was assessed by qPCR and western blot analysis.

---

**Fig S1.**
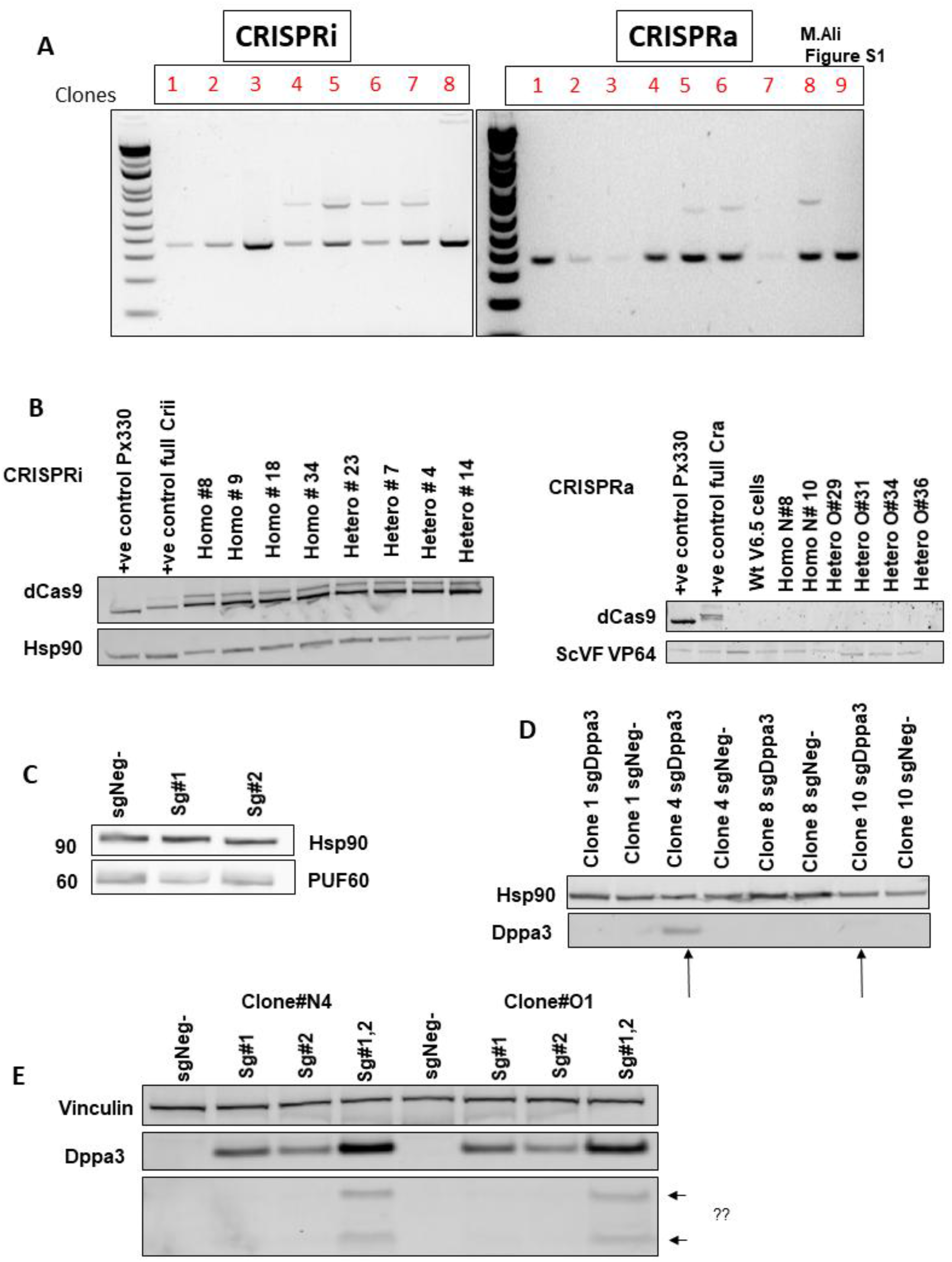
Generation and validation of CRISPRa and CRISPRi mESC lines. **A)** Genotyping of CRISPRa and CRISPRi cell lines using primers designed to amplify regions outside the insertion site. **B)** Western blot analysis detecting dCas9-KRAB expression in CRISPRi cells and dCas9-VP64 expression in CRISPRa cells. **C)** CRISPRa-mediated activation of *Puf60* expression in stable CRISPRa mESCs using control sgRNA, *Puf60* sgRNA1, and *Puf60* sgRNA2. **D)** Evaluation of CRISPRa efficiency across multiple clones following lentiviral transduction with Dppa3-targeting sgRNAs compared with sgRNA control cells. **E)** Assessment of transient plasmid-based sgRNA transfection efficiency in two independent CRISPRa clones using sgCtrl, sg1, sg2, and combined sg1/2 targeting constructs.

**Fig S2.**
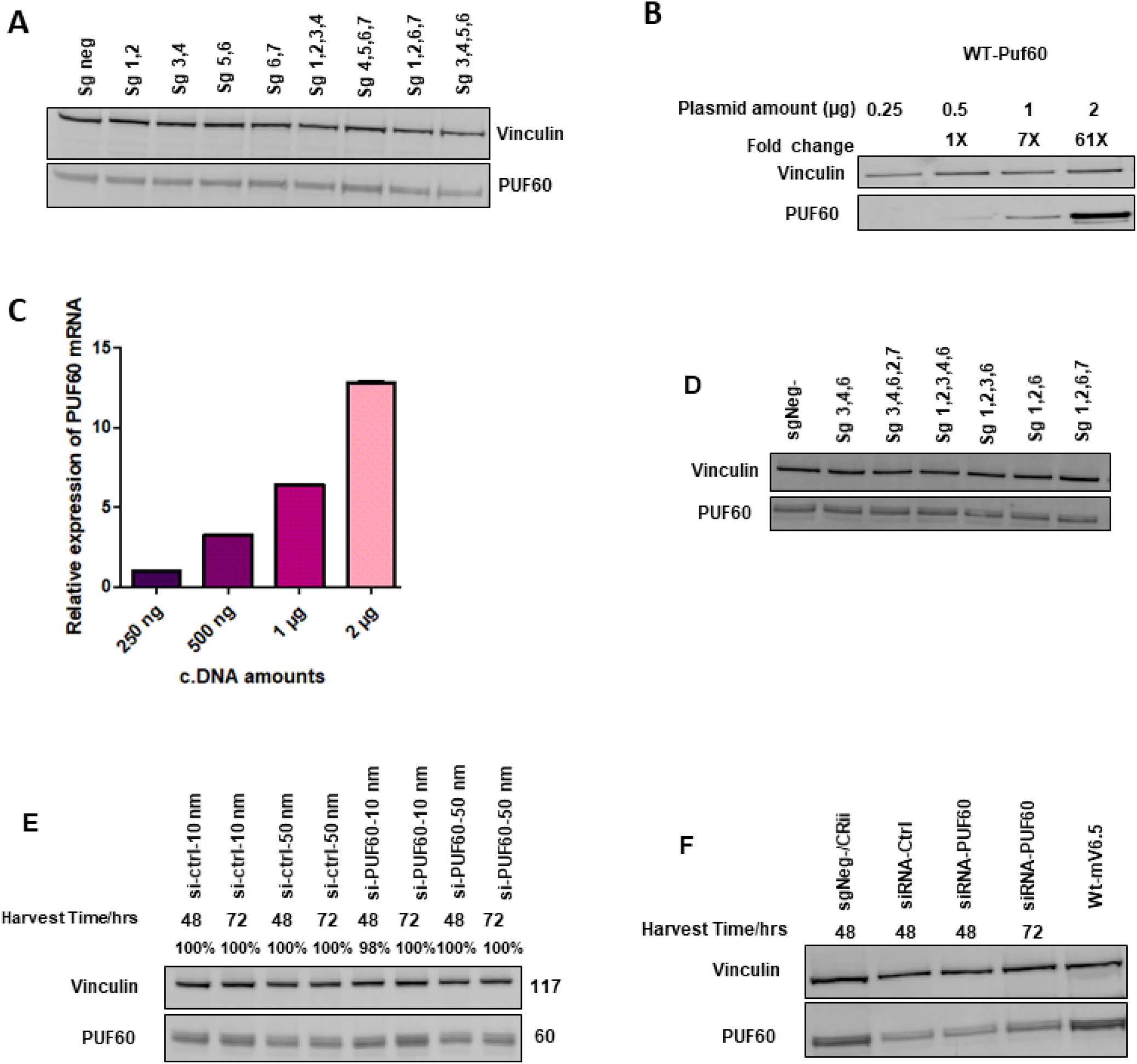
CRISPRi reveals resistance of Puf60 to transcriptional repression. **A)** Western blot analysis of CRISPRa cells transduced with control sgRNA or multiple combinations of *Puf60*-targeting sgRNAs 24 hours post-transduction. **B)** Western blot analysis of WT-Puf60 titration experiments. **C)** qPCR analysis demonstrating amplification of RNA expression with increasing cDNA amounts. **D)** Western blot analysis following targeting of Puf60 using multiple sgRNAs. **E)** Western blot analysis of siRNA-mediated knockdown of *Puf60*. **F)** Western blot analysis comparing siRNA-mediated and CRISPRi-mediated repression of Puf60.

